# Bitter taste receptor T2R14 and autophagy flux in gingival epithelial cells

**DOI:** 10.1101/2024.01.17.576143

**Authors:** Nisha Singh, Ben Ulmer, Manoj Medapati, Robert Schroth, Saeid Ghavami, Prashen Chelikani

## Abstract

**Macroautophagy (hereafter** autophagy) is a lysosomal degradation pathway that functions in nutrient recycling and as a mechanism of innate immunity. Previously we reported, a novel host-bacteria interaction between cariogenic *S. mutans* and bitter taste receptor (T2R14) in gingival epithelial cells (GEC) leading to an innate immune response. Further, *S. mutans* might be using the host immune system to inhibit other Gram-positive bacteria, such as *S. aureus*. To determine whether these bacteria exploit the autophagic machinery of GEC, it is first necessary to evaluate the role of T2R14 in modulating autophagic flux. So far, the the role of T2R14 in the regulation of autophagy is not well charcterized. Therefore, in this study, for the first time, we report that T2R14 downregulates autophagy flux in GECs and T2R14 knockout increases acidic vacuoles. Transmission electron microscopy morphometric results also suggested increased number of autophagic vesicles in T2R14 knockout GEC. Further, our results suggest that *S. mutans* competence stimulating peptide CSP-1 showed robust intracellular calcium release and this effect is both T2R14 and autophagy protein 7 dependent. In this study we provide the first evidence that T2R14 modulates autophagy flux in GEC. The results of current study culd have benefitial impact on the identifying the impact of T2R in regulation of immuno microenviroment of GEC and its impact in oral health.

## 1 Introduction

Autophagy is a lysosomal degradation pathway which plays a role in nutrient recycling and energy generation. It is also involved in the clearance of damaged proteins, organelles, and certain pathogens. Thus it might be involved as a part of the innate immune response [1, 2]. Previous studies have revealed that autophagy participates in odontoblast aging, tooth development, and many more physiological and pathological processes [3-5]. Involvement of many nutrient sensing G protein-coupled receptor (GPCR) like umami and fat taste receptors in the autophagy phenomenon is well understood [6, 7]. However, involvement of bitter taste receptors (T2Rs) in autophagy pathway is still unknown. To the best of our knowledge only one study has shown a direct link between T2Rs and autophagy in airway smooth muscles (ASM) cells. This study showed T2R agonists quinine and chloroquine alter mitochondrial function, induced cell death and changed the number of autophagosomes via autophagy protein 5 (ATG5) and the coiled-coil myosin-like BCL2-interacting protein 1 (Beclin 1) pathway in ASM cells [8, 9]. Chloroquine and quinine resulted in microtubule associated light chain protein (LC3β-II) accumulation and this LC3β-II accumulation in ASM cells was surprisingly reduced in the presence of the autophagy flux inhibitor bafilomycin A1 (Baf A1). Thus, this finding suggests T2R agonist-mediated cell death in human ASM cells is potentially regulated by changes in autophagy pathway [8].

Oral gingival epithelial cells (GECs) are first line of defense and produce antimicrobial peptides (AMPs) against various cariogenic bacteria. Invasion of non-phagocytic cells like epithelial, endothelial and fibroblast is a common strategy of eluding the host immune system for many pathogens. These pathogens developed various survival mechanisms inside the host cells and autophagy is one of them. Previous transcriptome analysis suggested significant expression of T2R14 in GEC compared to other GPCRs and Toll-like receptors [10]. In a follow up study, we suggested that T2R14 modulates Gram-positive bacterial internalization and survival in GECs [11]. The GECs infected with *Staphylococcus aureus* induced T2R14-dependent human β-defensin-2 (hBD-2) secretion, and T2R14 knockout affects the cytoskeletal reorganization, thereby inhibiting *S. aureus* internalization [11]. Since, the internalization of *S. aureus* is T2R14 dependent, it is imperative to elucidate whether *S. aureus* and other cariogenic bacteria *S. mutans* use autophagic machinery for their survival, and for evading the host immune system. In order to determine whether such bacteria exploit the autophagic machinery of GEC, it is first necessary to evaluate the role of T2R14 in modulating autophagic flux.

Autophagy is a fundamental cellular process relying on the formation of double membrane vesicles, called autophagosomes, for the delivery of cytoplasmic proteins and organelles to the lysosomal degradation machinery for energy production and cellular homeostasis [12, 13]. Number of autophagosomes are controlled via autophagy flux which is the outcome of the autophagosome formation and degradation [14]. The initiation and nucleation of autophagosome formation is controlled by two major inductive protein complexes, the mammalian target of rapamycin (mTOR) complex 1 (mTORC1) and the Beclin 1 complex. The activity of the mTOR-kinase suppresses autophagosome formation and acts as a negative regulator of autophagy. Rapamycin induces autophagic activity by inhibiting mTOR phosphorylation and thus preventing its kinase activity. Therefore, regulation of the autophagic flux under certain physiological or pathological conditions (e.g., an imbalance of nutrient supply, oral pathogen invasion) is of particular interest in GECs. In this study we provide evidence that T2R14 modulates autophagic flux in GECs, and that this phenomenon is ATG7 independent. Furthermore, our results also suggested that activation of T2R14 with *S. mutans* competence stimulating peptide (CSP-1) showed robust intracellular calcium release and this effect is both T2R14 and ATG7 dependent in GEC.

## 2 Materials and Methods

### 2.1 Reagents used in the study

Acridine orange (CAS Number: 65-61-2), bafilomycin A1 (Cat#B1793-10UG) and rapamycin were purchased from Sigma-Aldrich Co. Canada. Synthetic *S. mutans* competence stimulating peptide (CSP-1) and S. aureus auto inducer peptide (AIP) of 98% purity was purchased from GenScript (Piscataway, NJ, USA). Keratinocyte growth medium-2 (KGM-2) for OKF6 cell culture and Iscove’s modified Dulbecco’s medium (IMDM) for promyeloblast (HL-60) cells were purchased from promo cell (Heidelberg, Germany) and ATCC (Gaithersburg, MD), respectively. The following antibodies were purchased: mouse monoclonal anti-β-actin (#A5441) from Sigma Aldrich (Oakville, ON, Canada), goat anti-rabbit IgG-HRP conjugate (#17-6515) from Bio-Rad (Mississauga, ON, Canada), goat anti-mouse IgG-HRP conjugate (#A-10668) and rabbit polyclonal anti-T2R14 (#OSR00161W) from Thermo Fischer Scientific, Carlsbad, CA, USA, and rabbit polyclonal ATG7 from cell signalling.

### 2.2. Cell Line Used in the Study

The oral keratinocyte cell line OKF6 was a kind gift from Dr. Gill Diamond, University of Florida [15, 16]. The T2R14 KO and a non-targeting (MOCK) Alt^®^-R-control CRISPR-crRNA or MOCK OKF6 cells were generated by using the CRISPR-Cas9 technique as published in our previous study [10]. The promyelo-blast (HL-60) cells were cultured in IMDM supplemented with 20% FBS. The HL-60 cells were differentiated toward neutrophils using previously established protocols [17].

### 2.3. Bacterial Strains Used in the Study

The S. aureus (ATCC strain 6538, kind gift from Dr. Duan’s lab) and *S. mutans* strain UA159 was purchased from ATCC. The S. aureus strain is propagated in Luria-Bertani (LB) broth at 37 °C, under constant agitation. The *S. mutans* strain is propagated in Brain Heart Infusion (BHI) broth at 37 °C and 5% CO_2_, under constant agitation.

### 2.4 Co-culture assay on GEC

Briefly, for the co-culture assay, *S. aureus and S. mutans* were cultured overnight at 37°C with shaking at 200 rpm in their respective culture media and the GEC cells were infected with either *S. aureus* or *S. mutans* at a multiplicity of infection (MOI) of 50 and 100 for 16-18 hours respectively. Cell viability post-infection was assessed using the WST-1 assay[11]. After the treatments, conditioned medium (CM) was collected and filtered by using 0.2 μm nylon filter. The filtered CM was used to determine the dHL-60 cell migration assay as described previously [10].

### 2.5. Transmission Electron Microscopy Imaging of GECs

The OKF6 WT and T2R14 KO cells were fixed with 3% gluteraldehyde in 0.1 M Sorensen’s buffer for 3 h. After fixation the cells were suspended in sucrose solution and stored at 4 °C for processing. The above fixed cell suspensions were embedded into plastic resins and thin sections (90–100 nm) are placed on mesh copper grids. The copper grids are finally stained with osmium tetroxide and uranyl acetate [18]. The grids were imaged by using a Philips CM10 microscope at a magnification of 10,500x.

### 2.6 Acridine Orange Acidic Vacuole Assay

The autophagic flux was assessed using Acridine Orange (AO) vacuolar assay for detecting acidic vesicular organelles (AVO), which are the indispensable markers of autophagy [19, 20]. Acridine Orange staining induces green fluorescence in the cytosolic and nuclear fractions of cells whereas when it becomes protonated, emits red intense fluorescence upon fusion into the acidic environment, such as lysosomes. Thus, the ratio of red/green fluorescence intensity could be a suitable marker for AVO formation. GECs were seeded in 12 well plates under serum starved conditions and treated for 72 hours. After 72 hours, cells were stained with AO (final concentration of 1μg/ml) and incubated at 37°C for 10 mins in dark. After washing once with PBS, the flow cytometry analysis was performed for evaluating the autophagy flux (Red and Green fluorescence) using cytoflex flow cytometer [20, 21].

### 2.7 Immunoblot analysis

The protein expression level of ATG7 was quantified through immunoblotting. Cell lysates were prepared from control and ATG7 KD specific cells. Protein estimation in the samples was done using Bio-Rad DC protein estimation kit. Each sample, containing 20 μg of protein, was resolved on a 10% SDS-PAGE and subsequently transferred onto nitrocellulose membranes. These membranes were blocked using 5% non-fat dry milk for one hour at room temperature, followed by incubation with specific primary antibodies overnight at 4°C. The proteins were visualized by chemiluminescence, employing horseradish peroxidase (HRP)-conjugated secondary antibodies and enhanced chemiluminescence (ECL) substrate, with images captured on a ChemiDoc MP imaging system. Exposure times for chemiluminescent detection were set at 30 seconds for ATG7, and 2 seconds for the loading control, β-actin.

### 2.8 Intracellular calcium mobilization assay

Agonist induced intracellular calcium mobilization assays were performed as described previously[10, 22]. WT, T2R14KO, ATG7 KD and T2R14KO-ATG7KD cells (approx. 30,000/96 wells) were loaded with Fluo-4NW calcium dye and intracellular calcium response was measured after AIP-1 (50 μM), CSP-1(50 μM) and CSP-1 plus Baf A1 treatment. Intracellular calcium changes were quantitatively measured using a Flex Station® 3 MultiMode Microplate Reader, with results expressed as alterations in relative fluorescence units (RFU). Changes in intracellular concentration were recorded using Flex Station® 3 MultiMode Microplate Reader with data represented as change in relative fluorescence unit (RFU).

### 2.9 Immune cell (dHL-60) migration

dHL 60 cells were seeded on the top chamber of the CIM plate at a density of 4 × 10^5^ cells/well in HBSS containing 0.5% BSA. The bottom chamber was loaded with conditioned medium (CM) obtained from with or without *S. aureus* and *S. mutans* treatments in WT, T2R14KO, ATG7KD and T2R14KO-ATG7KD cells on the migration of differentiated HL-60 cells (dHL-60). The changes in cellular impedance or cell index (CI) were measured every 5 mins over 24 hours using RTCA software [10].

### 2.9 Knocking dowan of ATG7

Human GEC were initially placed in 12-well plates at a density of 5×104 cells per well and cultured in KGM-2 medium with growth supplements for 24 hours. When the cells reached approximately 40% confluency, they were exposed to 10 μg/ml of polybrene (Santa Cruz; sc-134,220) in KGM-2 basal medium for 1 hour. Following this, the cells were transfected with shRNA Lentiviral Particle encoding for shRNA ATG7 and scrambled control, which also carried the puromycin resistance marker (Santa Cruz; sc-41447-V). The transfection was conducted at 3 and 6 multiplicity of infections (MOI) for 12 hours, after which the medium was replenished for a 24-hour recovery period. To identify the cells that successfully incorporated the shRNA plasmids, they were grown in medium containing puromycin dihydrochloride (4 μg/ml) (Santa Cruz; sc-108071). The Western blot technique was then used to examine the ATG7 status of both the shRNA transfected cells and the scramble cells in isolated stable clones [23].

### 2.10. Statistical analysis

All experimental results are presented as the mean ± Standard error of the mean (SEM) from at least three independent experiments. GraphPad PRISM v9.0 (GraphPad Software, San Diego, CA, USA) was used to perform statistical significance analyses. One-way analysis of variance (ANOVA) was applied to compare more than three groups. Data presented as mean ± standard error of mean (SEM); *p<0.05, **p<0.01, ***p<0.001, ****p<0.0001 as indicated.

## 3 Results

### 3.1. T2R14 dependent autophagy flux in GEC

Our previous studies suggest T2R14 is the primary driver of innate immune responses to bacterial infection in GECs [10, 11]. To evaluate the effect of T2R14 in autophagy flux in GECs, oral keratinocyte cells, OKF6 Wt (WT) and OKF6 T2R14 KO (T2R14KO) cells were used as a model system. The autophagic flux was assessed using Acridine Orange vacuolar assay for detecting acidic vesicular organelles (AVO), which are the indispensable markers of autophagy [19, 20]. Both cell lines were serum starved for 72 hours to induce the autophagy [14, 24]. Cells were also treated with an autophagy flux inhibitor Bafilomycin A1 [Baf A1 (10 nM)] and an autophagy flux inducer, rapamycin [Rapa (1000 nM)] for 72 hours. Treatment of GECs with Baf A1 and Rapa for 72 hours did not lead to any significant change in cell viability (Fig 1A) and flow cytometry analysis of serum starved GECs did not result in significant cell death. After 72 hours, cells were stained with Acridine Orange (final concentration of 1μg/ml) and incubated at 37°C for 10 mins in dark. After washing once with PBS, the flow cytometry analysis was performed for evaluating the autophagy flux (Red and Green fluorescence) using cytoflex flow cytometer [20]. Acridine Orange staining induces green fluorescence in the cytosolic and nuclear fractions of cells whereas when it becomes protonated, emits red intense fluorescence upon fusion into the acidic environment, such as lysosomes. Our result suggested that serum starvation in T2R14KO cells leads to a significant increase in red fluorescence compared to WT (Fig. 1 B), suggesting more acidic vacuoles in T2R14KO cells. Further no change in the green fluorescence was observed (Fig. 1 C). Finally, the red/green fluorescence ratio were also significantly increased in T2R14KO serum starved condition as compared to WT serum starved group (Fig. 1 D). Further, treatments of WT and T2R14KO cells with autophagy flux inhibitor Baf A1 leads to complete loss off red fluorescence (Fig 1 E). Baf A1 treatments inhibits vacuolar H(+)-ATPase (V-ATPase), which results in the inability of the lysosome to acidify, thus no red fluorescence was observed. As expected, autophagy flux inducer Rapa leads to an increase in red fluorescence (Fig. 1 E). To corroborate the data obtained using Acridine Orange staining of AVO, transmission electron microscopy (TEM) was performed on WT and T2R14 KO GECs. TEM morphometric result suggested an increased number of autophagic vesicles (autophagosome and phagophore) in T2R14KO relative to the WT (Fig. 1 F-H). Autophagic vacuoles were classified as autophagosomes when they met the following criteria-1) Double membrane (complete or at least partially visible), 2) Absence of the ribosomes attached to the cytosolic side of the membrane [25].

**Figure 1.**
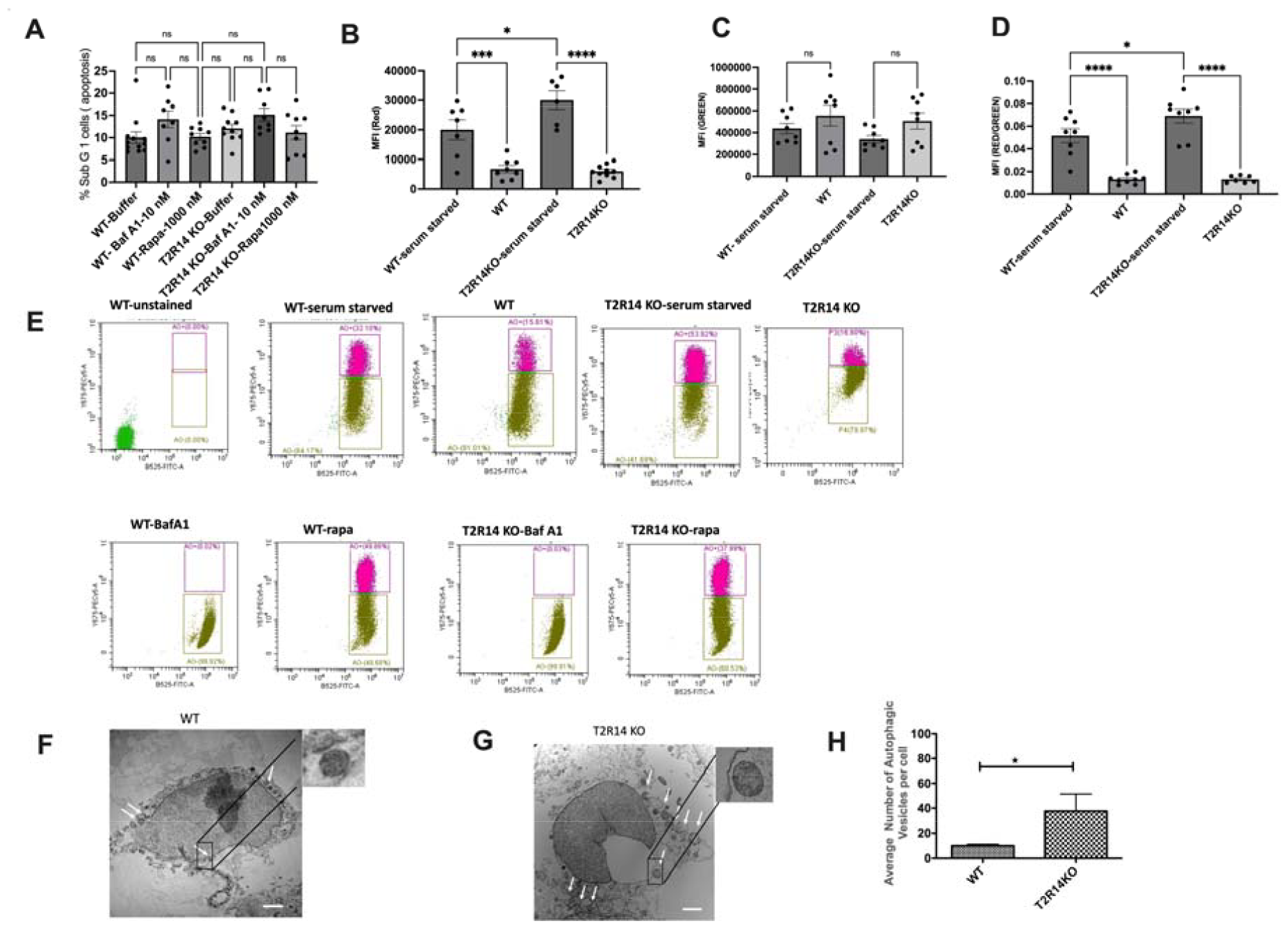
T2R14 dependent autophagy flux in gingival epithelial cells (GEC). **A. Effect of Baf A1 and Rapa on the GEC cell viability.** Flow cytometry cell-based assay suggested the effect of Baf A1(10 nM) and Rapa (1000 nM) on the cell growth in both WT and T2R14KO cells over the 72 hours. The bar plots represent SEM of 3 independent experiments performed in triplicates. **B-D. Acridine Orange staining of AVO for detecting autophagy flux in GEC**. Flow cytometry evaluation of red, green and red/green fluorescence ratio for detecting the autophagy flux in WT and T2R14KO in GEC under serum starved condition and in their respective normal growth condition. The bar graphs were generated using graph pad prism 9.0. Statistical significance was calculated using One-way ANOVA with Bonferroni’s post-hoc test *p<0.05, **p<0.01. The data represents SEM of 3-4 independent experiments. **E**. Acridine Orange staining of AVO for detecting autophagy flux in GEC. Representative raw traces showing flow cytometric detection of red and green fluorescence in acridine orange-stained GECs (WT and T2R14 KO) that were serum starved or treated with Baf A1 (10 nM) and Rapa (1000 nM) for 72 hours for detecting the autophagy flux. **F-H. Transmission electron microscopy (TEM) evaluation of autophagic vacuoles in WT and T2R14KO cells**. The white-headed arrow indicates the phagophore (incomplete autophagosome) and autophagosome double membrane structure (Insets) in respective GECs (**F and G**). Scale = 2 microns. **H**. Quantification of autophagosome based on the morphology (double membrane) in WT and T2R14 KO cells. Result represents mean value of autophagic vesicles per cell.

### 3.2 ATG7 independent autophagy flux in GEC

ATG7 is important for the formation of autophagic vesicles and directly involved in the processing of LC3-I to LC3-II [14, 26]. To check the involvement of ATG7 in autophagosome formation in GECs, ATG7 was knocked down in both WT and T2R14KO GECs using shRNA (Fig. 2 A). Loss of ATG7 in GECs resulted in non-significant changes of red fluorescence or ratio of red/green fluorescence (Fig. AI and AIII). However, we did not observe significant changes in red and red/green fluorescence after Acridine Orange staining in T2R14KO-ATG7KD (Fig 2B-E). This could be due to an ATG7-independent autophagy pathway in GECs. Previous studies suggested autophagosome formation and autophagy process is independent of ATG7 in different cell types [27, 28].

**Figure 2.**
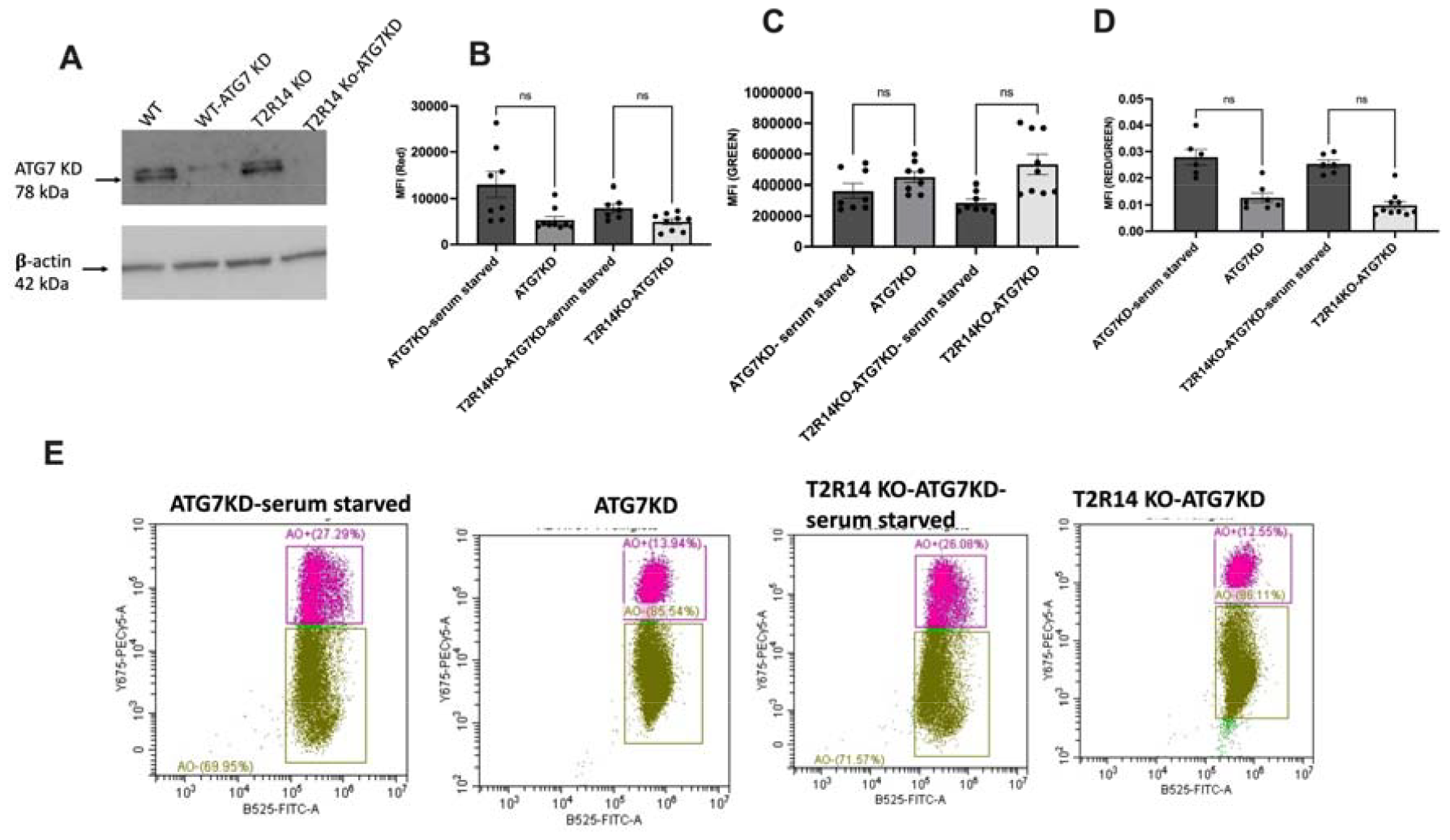
ATG7 independent autophagy flux in gingival epithelial cells (GEC). **A**. Western blot analysis of ATG7 knockdown in WT and T2R14KO GECs. ATG7 KD in WT and T2R14 KO GECs was done using shRNA lentiviral approach. **B-D. Acridine Orange staining of AVO for detecting autophagy flux in GEC**. Flow cytometry evaluation of red, green and red/green fluorescence ratio for detecting the autophagy flux in ATG7Kd and T2R14KO-ATG7KD in GEC under serum starved condition and in their respective normal growth condition. The bar graphs were generated using graph pad prism 9.0. Statistical significance was calculated using One-way ANOVA with Bonferroni’s post-hoc test *p<0.05, **p<0.01. The data represents SEM of 3-4 independent experiments. **E**. Acridine Orange staining of AVO for detecting autophagy flux in GEC. Representative raw traces showing flow cytometric detection of red and green fluorescence in acridine orange-stained GECs (ATG7KD T2R14-ATG7KD) that were serum starved for 72 hours for detecting the autophagy flux.

### 3.3 Intracellular calcium mobilization assay in GEC

There have been several reports suggesting involvement of calcium in the autophagy pathway [29, 30]. Calcium is known to act as both a promoter and inhibitor of autophagy under different circumstances. It is well known that activation of T2Rs leads to intracellular calcium release via T2Rs-G_βγ_ –PLC_β_-IP_3_-Ca^2+ [10, 31]^. To elucidate whether changes in autophagy flux could affect intracellular mobilization of Calcium ion via activation of endogenous T2R14, GEC cells were stimulated with Baf A1(10nM) and rapa (1000nM) and intracellular calcium release were measured. BafA1 leads to no significant changes in intracellular calcium release in ATG7 KD or T2R14KO cells compared to WT treated cells (Fig. 3A). Our previous findings suggested that in GECs, competence stimulating peptide 1 (CSP-1) from cariogenic bacteria *S. mutans* exhibited T2R14 dependent intracellular calcium release [10]. We also demonstrated T2R14 dependent *S. aureus* bacterial internalization and release of antimicrobial peptide (HBD2) in GEC [11]. However, the effect of autoinducing peptide 1 (AIP-1) from *S. aureus* and CSP-1 from *S. mutans* on intracellular calcium release in ATG7KD cells was not known. ATG7KD and T2R14KO GECs showed lower calcium release as compared to the WT upon CSP-1 treatment (Fig. 3 B). Next, the combined effect of Baf A1 and CSP-1 on calcium signalling was analyzed. Result suggested there was no change in intracellular calcium release upon co-treatment of GECs with Baf A1 and CSP-1 compared to CSP-1 alone treatment (Fig 3 C). This suggests that T2R14 mediated calcium release was not involved in T2R14-dependent autophagy flux in GEC.

**Figure 3.**
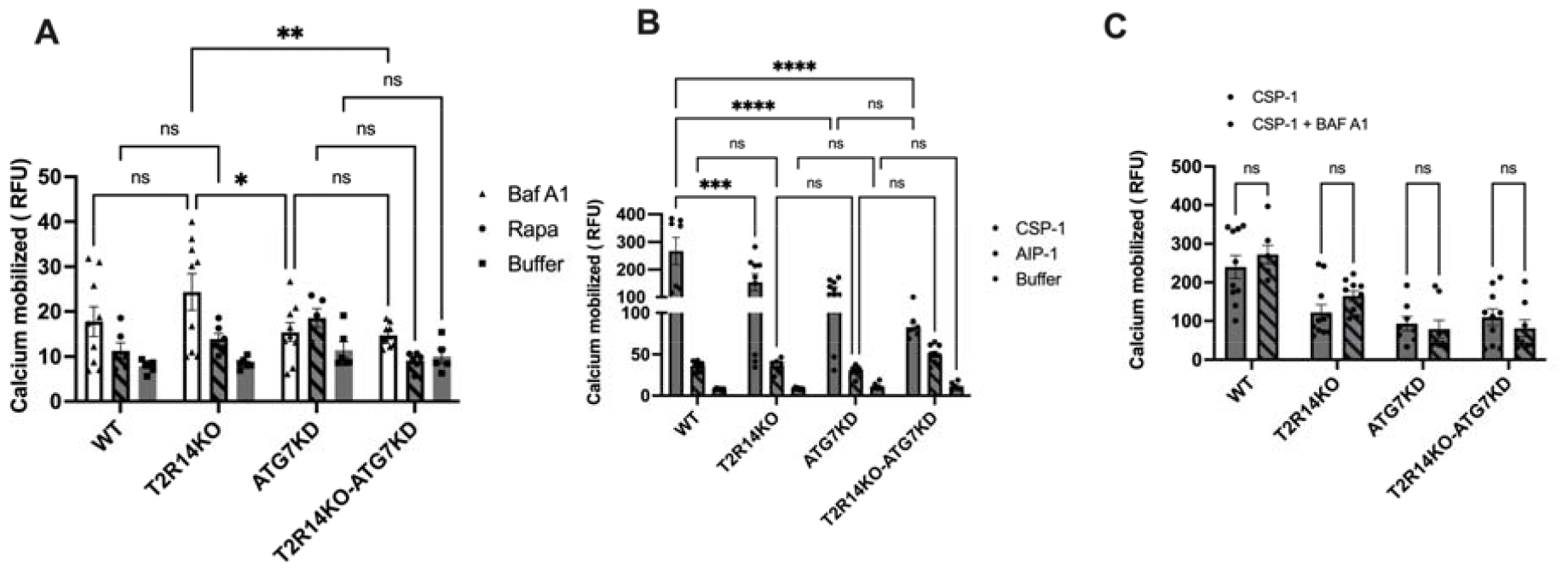
Intracellular calcium mobilization assay in GEC. A. Pharmacological characterization of autophagy flux inhibitor, Baf A1 (10 nM) and autophagy inducer rapa (1000 nM) on the calcium mobilization in GEC. WT, T2R14KO, ATG7 KD and T2R14KO-ATG7KD cells (approx. 30,000/96 wells) were loaded with Fluo-4NW calcium dye and intracel cellular calcium dye and intracell calcium response was measured after Baf A1 and rapa treatment. Data shown are mean ± SEM from three independent experiments done in triplicate. **B. Effect of *S. aureus* auto inducer peptide (AIP) and *S. mutans* competence stimulating peptide (CSP-1) on calcium mobilization in GEC cells**. Intracellular calcium release measurement after application of autoinducer peptides (AIP from *S. aureus* and CSP-1 from *S. mutans* 50μM each) in GEC cells. Results are from three independent experiments done in triplicate. **C**. Combined effect of Baf A1 and CSP-1 on calcium signalling in GEC. WT, T2R14KO, ATG7 KD and T2R14KO-ATG7KD cells (approx. 30,000/96 wells) were loaded with Fluo-4NW calcium dye and intracellular calcium response was measured after CSP-1 and CSP-1 plus Baf A1 treatment. Data shown are mean ± SEM from three independent experiments done in triplicate.

### 3.4. Immune cell (dHL-60) migration

Neutrophils are the first immune cells recruited to the site of gingival tissue inflammation upon bacterial infection. Our previous result suggested that treatment of GECs with synthetic CSP-1 led to an increased secretion of the antimicrobial peptides (AMPs), chemokine interleukin 8 (IL-8) and human beta defensin 2 (HBD-2), in a T2R14-dependant manner [10]. To investigate whether the oral bacteria infected GECs recruit immune cells that is T2R14 and/or ATG7 dependent, we tested effect of conditioned medium (CM) obtained from *S. aureus* and *S. mutans* treated GECs on the migration of neutrophil like cell line differentiated HL-60 cells (dHL-60). Treatment of dHL-60 cells with the aforementioned conditioned media led to no changes in dHL-60 cells, which is consistent with our previous report [10]. Previously we observed a significant decrease in dHL-60 migration upon treatment with conditioned media from T2R14KO cells primed with synthetic CSP-1 as compared to mock cells. We did not observe a similar effect on dHL-60 migration when treated with conditioned media from either *S. aureus* or *S. mutans* (Fig 4 A-B). This could be due to multiple reasons including the low levels of peptides secreted by the microbes in our assay system, internalization of microbes by GECs, the differences in post-translational modification of the peptides secreted by the microbes versus synthesized peptides. Further, treatment of dHL-60 cells with BafA1 and Rapa were performed. Result suggested there was no effect of these compounds on migration of neutrophil like cells (Fig. 4 C).

**Figure 4.**
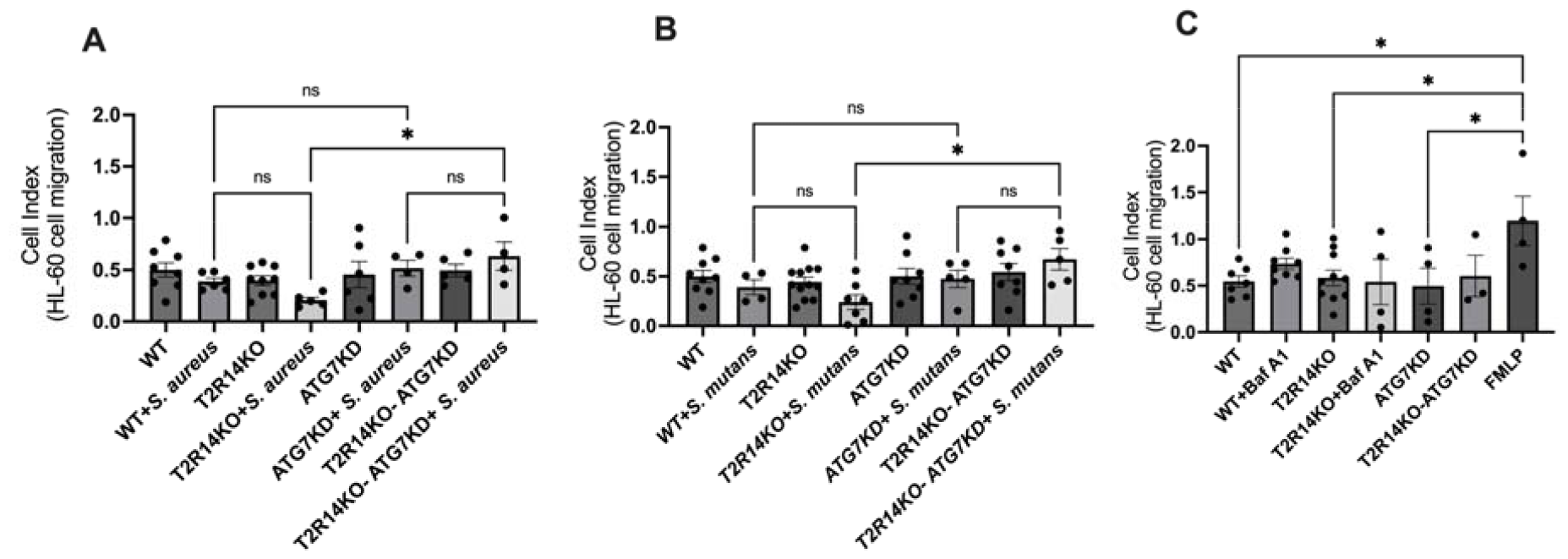
Immune cell (dHL-60) migration. **A-B**. Conditioned medium (CM) obtained from with or without *S. aureus* and *S. mutans* treatments in WT, T2R14KO, ATG7KD and T2R14KO-ATG7KD cells on the migration of differentiated HL-60 cells (dHL-60). The changes in cellular impedance or cell index (CI) were measured every 5 mins over 24 hours using RTCA software. The bar graph shows change in the migration of dHL-60 cells over a period of 24 hours. The bar plots represent SEM of 3 independent experiments and were generated using graph pad prism 9.0. Statistical significance was calculated using One-way ANOVA with Bonferroni’s post-hoc test *p<0.05, **p<0.01. **C**. Effect of conditioned medium (CM) obtained from with or without BafA1 (10 nM) treatment in WT and T2R14KO and from ATG7KD and T2R14KO-ATG7KD cells on the migration of differentiated HL-60 cells (dHL-60). The changes in cellular impedance or cell index (CI) were measured every 5 mins over 24 hours using RTCA software. FMLP was included as a positive control for neutrophil chemoattractant. The bar plots represent SEM of 2-3 independent experiments performed in triplicates. *P<0.05 showed statistically significant.

## 4 Discussion

Autophagy is a physiological compensation process involved in the maintenance of cell homeostasis via removing misfolded proteins and damaged organelles. It is also involved in regulation of host cell response against microbial infection. However, the mechanism by which these oral pathogens utilize autophagy as a survival mechanism and elude the host immune system is not well understood. In this study, we characterize the role of T2R14 in modulating autophagy flux in GECs. On the basis of Acridine Orange staining of AVO (Fig. 1), the presence of T2R14 in GECs increases the number of acidic active lysosomes. This could be due to an effect of T2R14 on the activity of lysosomal enzymes, or an effect of T2R14 on the intracellular pH of lysosomes. Further study is required to confirm the effects of T2R14 on the lysosomal activity and on the intracellular pH of lysosomes.

Both *ATG5* and *ATG7* are believed to be indispensable for autophagy pathway [32]. Thus, in the present study we knock down the *ATG7* gene for performing the autophagy flux. Our result (Fig. 2) suggested that loss of ATG7 in GECs resulted in non-significant changes of red fluorescence or ratio of red/green fluorescence. This could be due to an ATG7-independent autophagy pathway in GECs. Previous studies also suggested autophagosome formation and autophagy process is independent of ATG7 in different cell types [27, 28]. The presence of ATG5/ATG7-independent, alternative autophagy pathway has been confirmed in a variety of cells which includes fibroblasts and preadipocytes obtained from *Atg5*^flox/flox^ mice [33], thymocytes obtained from an *Atg5*^−/−^ mice embryo [28], mouse pancreatic β-cells [34] and induced pluripotent stem cells (iPSCs) [33].

Previous reports suggested the involvement of calcium in the autophagy pathway [29]. Calcium is known to act as both a promoter and inhibitor of autophagy under different circumstances. It is well established that activation of T2Rs leads to intracellular calcium release [10, 31]. Our results suggested there was no change in intracellular calcium release upon co-treatment of GECs with Baf A1 and CSP-1 compared to CSP-1 alone treatment (Fig 3C). This suggests that T2R14 mediated calcium release was not involved in T2R14-dependent autophagy flux in GEC. Studies suggested that intracellular calcium is not always associated with increase or decrease in autophagy flux [30, 35]. In general, the final effect of calcium on autophagy depends on spatiotemporal characteristics and the amplitude of calcium signals as well as on cell growth conditions (e.g., nutrient and growth factor availability) and type of autophagy [36].

The infiltration of immune cells such as neutrophils and macrophages lead to gingival tissue inflammation [37]. In recent years, autophagy machinery has emerged as a central regulator of innate immune functions, cytokine production, immune cells differentiations and pathogen clearance [38]. Neutrophils are the first line of immune cells recruited to the site of gingival tissue inflammation upon bacterial infection. Thus, the effect of conditioned medium obtained from *S. aureus* and *S. mutans* treated GECs on the neutrophil like cell line dHL-60 was investigated. Treatment of dHL-60 cells with conditioned media led to no changes in dHL-60 cells, which is consistent with our previous report [10]. Previously we observed a significant decrease in dHL-60 migration upon treatment with conditioned media from T2R14KO cells primed with synthetic CSP-1 as compared to mock cells. We did not observe a similar effect on dHL-60 migration when treated with conditioned media from either *S. aureus* or *S. mutans* (Fig 4 A-B). It is possible that S. aureus and S. mutans once internalized inside the GECs are unable to elicit sufficient cytokine production involved in the neutrophil migration. Since, T2R14 differentially modulates the internalization of *S. aureus* and *S. mutans* inside GECs, thus it is imperative to analyze how these gram-positive bacteria manipulate host cells for their survival and their effects on oral innate immunity. Further studies to investigate whether cariogenic and non-cariogenic bacteria use similar autophagy process to form their replicative niche in GECs is warranted.

Previous study suggested that the autophagosome or autophagic vesicles formation involves both non-canonical and canonical pathways [39]. Future studies evaluating the role of lysosomal small Rho GTPase, mTOR and Beclin 1 complexes in the regulation of T2R mediated autophagy in GECs and other cell types can be pursued. It is far beyond the aims of the present study to investigate the complete signaling pathway involved in the autophagic machinery in the context of oral innate immunity. On the other hand, autophagy could be a master regulator of chemokine and cytokine secretion from the mammalian cells through secretory autophagy and unfolded protein response [40, 41]. Future investigations addressing if T2Rs are invovled in regulation of secretion and processing of GEC originated chemokines and cytokines via secretory autophagy and unfolded protein response is very much needed.

## Author Contributions

N.S; B.U and M.M.R conducted the experiments and analyzed the data. N.S and P.C wrote the manuscript. P.C; N.S and S.G contributed to the conceptualization, design and data interpretation. All authors critically revised the manuscript.

## Funding

This work was supported by Grant No. PJT-159731 from the Canadian Institute of Health Research (CIHR) to PC. S.G., was supported by the CancerCare Manitoba Operating grant (763117252) funded by CancerCare Manitoba Foundation.

## Institutional Review Board Statement

Not applicable.

## Informed Consent Statement

Not applicable.

## Data Availability Statement

The data that support the findings of this study are available on reasonable request from the corresponding author.

## Acknowledgments

We thank Dr. Ryan Cunnington for critically reviewing the manuscript and generous support of Dr. Christine Zhang, flow cytometry core facility manager for the Acridine Orange staining data acquisition by flow cytometry.

## Conflicts of Interest

The authors declare no competing financial interests.

## Notes

### Competing Interest Statement

The authors have declared no competing interest.

